# Microcin MccI47 selectively inhibits enteric bacteria and reduces carbapenem-resistant *Klebsiella pneumoniae* colonization *in vivo* when administered *via* an engineered live biotherapeutic

**DOI:** 10.1101/2021.12.17.473159

**Authors:** Benedikt M. Mortzfeld, Jacob D. Palmer, Shakti K. Bhattarai, Haley L. Dupre, Regino Mercado-Lubo, Mark W. Silby, Corinna Bang, Beth A. McCormick, Vanni Bucci

**Affiliations:** Department of Microbiology and Physiological Systems, University of Massachusetts Medical School, Worcester, MA, USA; Program in Microbiome Dynamics, University of Massachusetts Medical School, Worcester, MA, USA; Department of Zoology, University of Oxford, Oxford, United Kingdom; Department of Biochemistry, University of Oxford, Oxford, United Kingdom; Department of Bioengineering, University of Massachusetts Dartmouth, North Dartmouth, MA, USA; Department of Biology, University of Massachusetts Dartmouth, Dartmouth MA, USA; Institute of Clinical Molecular Biology, Christian-Albrechts-Universität zu Kiel, Kiel, Germany; Program in Systems Biology, University of Massachusetts Medical School, Worcester, MA, USA

## Abstract

**Background:** The gastrointestinal (GI) tract is the reservoir for multidrug-resistant (MDR) pathogens, specifically carbapenem-resistant (CR) *Klebsiella pneumoniae* and other *Enterobacteriaceae*, which often lead to the spread of antimicrobial resistance genes, severe extraintestinal infections, and lethal outcomes. Selective GI decolonization has been proposed as a new strategy for preventing transmission to other body sites and minimizing spreading to susceptible individuals.

**Results:** Here, we purify the to-date uncharacterized class IIb microcin I47 (MccI47) and demonstrate potent inhibition of numerous *Enterobacteriaceae*, including multidrug-resistant clinical isolates, *in vitro* at concentrations resembling those of commonly prescribed antibiotics. We then genetically modify the probiotic bacterium *Escherichia coli* Nissle 1917 (*EcN*) to produce MccI47 from a stable multicopy plasmid by using MccI47 toxin production in a counterselection mechanism to engineer one of the native *EcN* plasmids, which renders provisions for inducible expression and plasmid selection unnecessary. We then test the clinical relevance of the MccI47-producing engineered *EcN* in a murine CR *K. pneumoniae* colonization model and demonstrate significant MccI47-dependent reduction of CR *K. pneumoniae* abundance after seven days of daily oral live biotherapeutic administration without disruption of the resident microbiota.

**Conclusions:** This study provides the first demonstration of MccI47 as a potent antimicrobial against certain *Enterobacteriaceae*, and its ability to significantly reduce the abundance of CR *K. pneumoniae* in a preclinical animal model, when delivered from an engineered live biotherapeutic product. This study serves as the foundational step towards the use of engineered live biotherapeutic products aimed at the selective removal of MDR pathogens from the GI tract

## Background

Modern health care is challenged by the increasing emergence of multi-drug resistant (MDR) pathogens, including carbapenem-resistant (CR) *Enterobacteriaceae*, which are responsible for millions of infections, tens of thousands of deaths, and billions of dollars in health care costs every year[1-3]. In 2017 the World Health Organization (WHO) released a list of bacteria, for which new antibiotics are urgently needed, and considered CR *Enterobacteriaceae* including *Klebsiella pneumoniae, Escherichia coli, Salmonella enterica*, or *Enterobacter cloaceae* of the highest priority[4]. While *K. pneumoniae* is not known to cause gastrointestinal (GI) disease, its stable colonization of the intestine is the main reservoir for infections and transmission[5]. MDR *Klebsiella* species are responsible for outbreaks in nursing homes and long-term care facilities[6] and for spreading antibiotic resistance among other members of the GI microbiota[7, 8]. Moreover, GI colonization by certain *K. pneumoniae* contributes to the development of inflammatory bowel disease[9] and fatty acid liver disease[10], making its selective removal a desireable milestone in modern medicine[11]. However, the paucity of treatment options for the elimination of CR *K. pneumoniae* begs for the development of new strategies to eradicate it from the GI tract.

The human gut microbiome is a densely populated ecosystem, where our bacterial symbionts are in constant competition for survival[12]. A major strategy employed by bacteria to outcompete their neighbors is the production of highly specific antimicrobial peptides, so called bacteriocins, that target closely related species or strains[13]. While small molecule antimicrobials, such as penicillin[14], have been exploited for decades as traditional antibiotics, antimicrobial peptides as therapeutic agents have just recently gained widespread interest as potential treatments of MDR pathogens[15, 16]. Siderophore antimicrobial peptides (sAMPs), including class IIb microcins, are ribosomally synthesized and post-translationally modified peptides (RiPPs) consisting of an antimicrobial peptide (AMP) linked to an iron-chelating molecule, a siderophore (*e*.*g*., enterobactin)[17]. Several studies have pointed to class IIb microcins as possible players capable of modulating the colonization dynamics of pathogenic *Enterobacteriaceae* in the GI tract[18-21], however, to date only three of them (MccM, MccH47, MccE492) have been characterized in detail[22-25]. MccI47 from *E. coli* strains H47 and CA46 has been identified based on sequence homology to known sAMPs, but it has never been overexpressed heterologously, purified, and no data demonstrating antimicrobial activity have been presented before[26]. In this study we first leverage our recently established pipeline for sAMPs purification[20, 21] to provide the first ever characterization of MccI47 as a potent antimicrobial against multiple *Enterobacteriaceae*. We then perform rational genetic manipulation of the probiotic *E. coli* Nissle 1917 (EcN) by engineering one of its two native plasmids to produce MccI47 without the need of any selection and demonstrate the ability of this new EcN strain in reducing CR *K. pneumoniae* abundance *in vivo*.

## Results

To expand the library of potent antimicrobials against MDR *Enterobacteriaceae*, we overexpressed MccI47 heterologously in *E. coli* and tested its inhibitory activity against a collection of enteric bacteria, including MDR clinical isolates. Specifically, we created a plasmid that contained the structural microcin (*mciA*) and immunity (*mciI*) gene under arabinose induction, followed by the genes necessary for posttranslational modification with monoglycosylated enterobactin (MGE) *mchCDEFA* (as in[20]) (**Figure 1A**). Highlighting the differences in activity spectrum and efficacy, we directly compared the potency of MccI47 with MccH47 (**Figure S1**), a well-characterized sAMP that has been used in previous proof-of-concept engineered biotherapeutics against *Salmonella*[20, 21]. Using static inhibition assays, we demonstrate that both microcins have strong inhibitory activity against extended-spectrum beta-lactamase (ESBL) producing *E. coli* (BAA 196). Intriguingly, MccI47 also inhibits the growth of CR *K. pneumoniae* (BAA 1705), while MccH47 has no effect (**Figure 1B**). We purified MccI47 with a maltose-binding protein (MBP)-microcin fusion protein[20] and direct comparison of MccH47 and MccI47 in equal quantities reproduced the results obtained using the live-producing bacteria, highlighting MccI47’s efficacy against CR *K. pneumoniae* (BAA 1705) (**Figure 1C**). We next assessed the potency of MccI47 in minimum inhibitory concentration (MIC) assays against a panel of 20 different target strains (**Table 1**). Overall, MccI47’s potency is comparable to that of commonly prescribed broad-spectrum antibiotics, while being highly specific for members of the *Enterobacteriaceae* family[27]. Compared to MccH47, MccI47 is approximately three-times more effective against various *E. coli* strains (0.2-0.6 µM), four-times more effective against *Shigella flexneri* (0.06 µM), and seven-times more effective against *Salmonella* Typhimurium (0.9-1.7 µM). More importantly, MccI47 but not MccH47 demonstrates activity against *Enterobacter cloacae* (19.5 µM), *Serratia marcescens* (15.1 µM), and *Klebsiella pneumoniae* (2.0-5.1 µM). Interestingly, neither of the two toxins showed activity against *Proteus mirabilis, Klebsiella oxytoca* or any tested bacteria outside of the *Enterobacteriaceae* family. We did not detect any difference in potency in targets carrying MDR genes, which is likely because the mode of entry of MccI47 into the target cells is independent of the resistance mechanisms of commonly used antibiotics[28]. Taken together these data demonstrate the discovery of MccI47 as a new potent AMP with selective activity against several *Enterobacteriaceae* members, including a strong inhibitory effect against CR *K. pneumoniae* representatives.

**Table 1:** Results of Minimum Inhibitory Concentration (MIC) Assays of purified MccI47 against bacterial isolates, including MDR *Enterobacteriaceae* in comparison to MccH47[20]. Asterisks indicate MDR strains.

| **Bacterial species** | **Strain** | **Origin** | **<u>MccI47</u>** |  | **<u>MccH47[20]</u>** |
| --- | --- | --- | --- | --- | --- |
|  |  |  | **<u>MIC (µg/ml)</u>** | **<u>MIC (µM)</u>** | **<u>MIC (µM)</u>** |
| *Acinetobacter baumannii* * | BAA-1790 | Human clinical isolate (sputum) | >138.8 | >19.6 | >113 |
| *Enterobacter cloacae* * | BAA-2341 | Human clinical isolate | 138.4 | 19.5 | >113 |
| *Escherichia coli* | 25922 | Human clinical isolate | 1.6 | 0.2 | 5.3 |
| *Escherichia coli* * | BAA-196 | Human clinical isolate | 4.4 | 0.6 | 1.8 |
| *Escherichia coli* | DH5α | Laboratory | 2.3 | 0.3 | 1.1 |
| *Klebsiella oxytoca* * | 51983 | Human clinical isolate (blood) | >197.3 | >27.8 | >113 |
| *Klebsiella oxytoca* | 700324 | Human isolate (bioMérieux, Inc.) | >197.3 | >27.8 | >113 |
| *Klebsiella pneumoniae* * | BAA-1705 | Human clinical isolate (urine) | 36.1 | 5.1 | >113 |
| *Klebsiella pneumoniae* * | BAA-2146 | Human clinical isolate | 29.4 | 4.2 | >113 |
| *Klebsiella pneumoniae* * | BAA-2342 | Human clinical isolate | 16.4 | 2.3 | >113 |
| *Klebsiella pneumoniae* * | BAA-2524 | Human clinical isolate | 14.7 | 2.0 | >113 |
| *Proteus mirabilis* | 29906 | Human clinical isolate (urogenital) | >197.3 | >27.8 | 5.3 |
| *Pseudomonas aeruginosa* | PA14 | Human clinical isolate | >138.8 | >19.6 | >113 |
| *Salmonella* Typhimurium | 19585 | Derived from LT2 (natural source) | 8.0 | 1.1 | 8.6 |
| *Salmonella* Typhimurium | 29630 | Derived from LT2 (natural source) | 6.3 | 0.9 | 6.3 |
| *Salmonella* Typhimurium* | BAA-190 | Human clinical isolate | 12.6 | 1.7 | 12.7 |
| *Serratia marcescens* | DB11 | Derived from DB10 (*Drosophila*) | 107.0 | 15.1 | >113 |
| *Shigella flexneri* | 2457T | Human clinical isolate | 0.4 | 0.06 | 2.4 |
| *Shigella flexneri* | M90T | Human clinical isolate | 0.4 | 0.06 | 4.4 |
| *Staphylococcus aureus* | 27661 | Human isolate | >197.3 | >27.8 | >113 |

**Figure 1:**
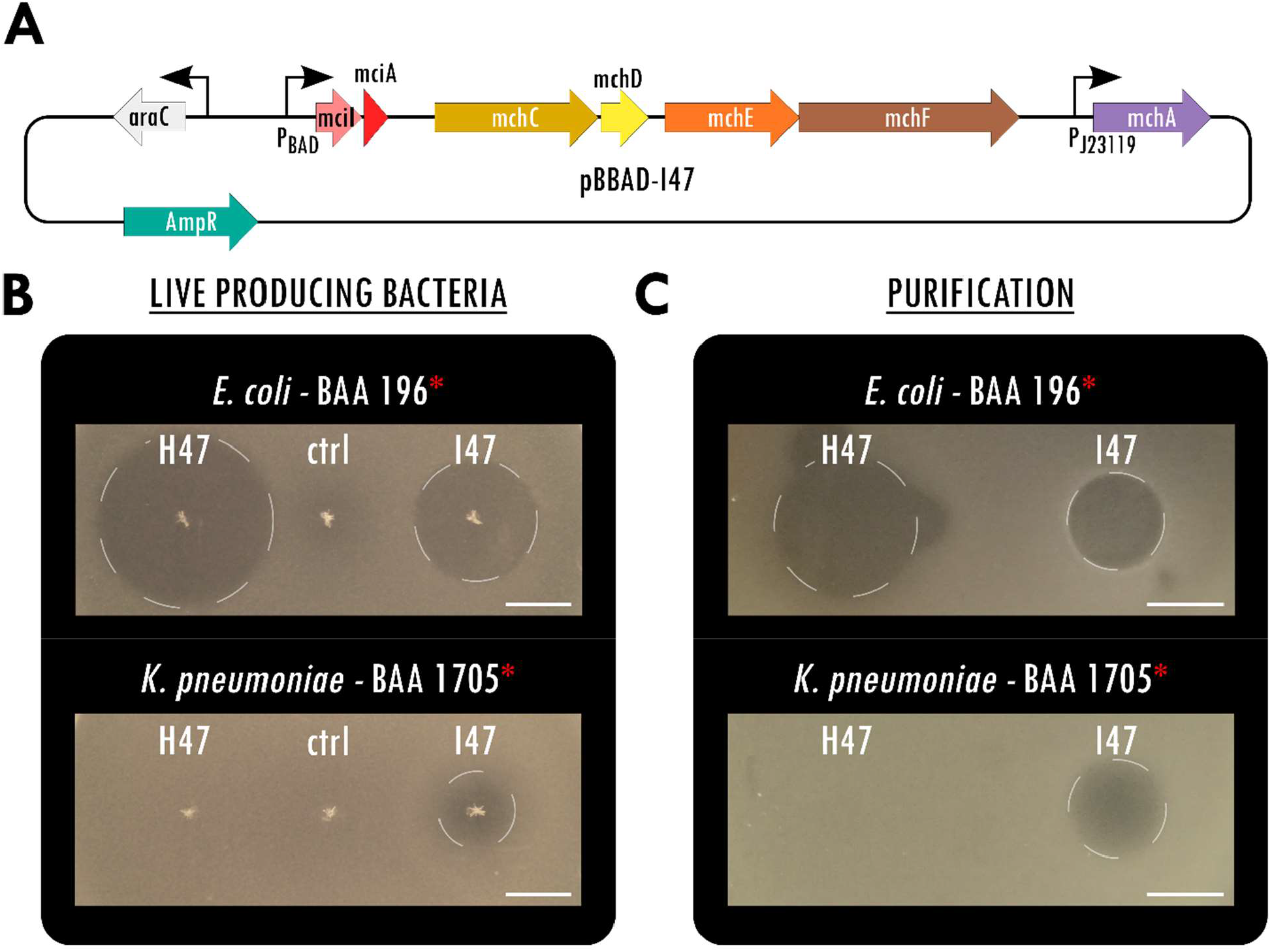
MccI47 is a potent inhibitor of MDR *Klebsiella pneumoniae*. (A) Plasmid map of pBBAD-I47, a pUC19-based plasmid producing MccI47-MGE from *mciA*, the immunity peptide *mciI* as well as the genes needed for post-translational modification *mchCDEFA*. (B) Static inhibition assays comparing MccH47 and MccI47 activity from single colony production against multidrug-resistant *Escherichia coli* (top) and *Klebsiella pneumoniae* (bottom). The control strain (ctrl) harbors the same backbone plasmid (pUC19) without microcin production. (C) Static inhibition assays comparing purified MccH47 and MccI47 activity against multidrug-resistant *Escherichia coli* (top) and *Klebsiella pneumoniae* (bottom). After purification, 3.5 µg of each microcin were dried on the media before the overlay with the target bacteria. Scale bars: 1 cm.

Due to the potent inhibitory effect against *K. pneumoniae* observed *in vitro*, we hypothesized that MccI47 could be used as a novel narrow-spectrum antimicrobial to reduce CR *K. pneumoniae* colonization in the GI tract in a murine model. We therefore developed an engineered *EcN* strain with MccI47-production from a native, modified multicopy plasmid, allowing for long-term plasmid retention without the need of selection. We chose *EcN* as vehicle strain because it has been used in its wildtype form as a live probiotic supplement for several human diseases for over a century[29], carries genomic islands that promote anti-inflammatory responses[30], and was found to be highly refractory to exogenous plasmid acquisition[31]. *EcN* is used for the construction of genetically engineered bacteriotherapeutic agents which are being tested in multiple ongoing clinical trials[32]. While recent studies relied on chromosomal integration[33, 34], we sought to maximize MccI47 output using a multicopy plasmid[21, 35, 36] and developed an expression system using the native, self-retaining *EcN* plasmid pMut2 as backbone for constitutive heterologous protein expression[36]. We first cured *EcN* from pMut2 by utilizing MccI47 (*mciA*) in a counterselection approach (**Figure 2A**). For the curing plasmid pCure2-I47 (**Figure S2A**) we employed pMut2 as backbone for plasmid competition, while harboring a chloramphenicol resistance, and placing *mciA* without its signal peptide for secretion and the protective immunity gene under isopropyl β-d-1-thiogalactopyranoside (IPTG) induction. Strong antibiotic selection allowed us to first displace pMut2, before inducing *mciA* expression and thus cell death in cells harboring pCure2-I47 (**Figure 2A**). The resulting cells cured of pMut2 and pCure2-I47 (**Figure S2B, Table S1**) were then transformed with the MccI47-producing plasmid (pMut2-I47) for *in vivo* applicability (**Figure 2B**). pMut2-I47 contains the same genetic machinery for MccI47 production and export as pBBAD-I47 (**Figure 1A**), however, it uses a proD-like[37] insulated strong constitutive promoter (J23119) for *mciA/mciI* expression for maximum microcin output *in vivo*. Further, the pMut2 backbone allows plasmid retention without selection pressure. Since wildtype *EcN* harbors genes for the microcins MccM (*mcmIA*) and MccH47 (*mchIB*), we generated an *EcN* MccH47/MccM knockout strain (*EcN*^*ΔH ΔM*^) to exclude potential inference by these microcins in the downstream experiments (**Figure S3, Figure S4**). Therefore, the following strains were used in the remaining set of experiments: (i) *EcN*, (ii) *EcN*^*ΔH ΔM*^, and (iii) *EcN*^*ΔH ΔM*^*-I47*, with the latter harboring pMut2-I47. Since pMut2-I47 is 2.6-times larger than the native pMut2 (5514 bp), we evaluated pMut2-I47’s retention in a serial passage experiment without antibiotic selection for ten consecutive days and approximately 200 generations in total. We did not observe a reduction in colony forming units (CFUs) carrying the plasmid (linear mixed effect modeling, p value > 0.05; **Figure 2C**) but intriguingly record a higher plasmid copy number for *EcN*^*ΔH ΔM*^*-I47* (mean ± SD, 17.4 ± 1.5) compared to *EcN* (12.9 ± 0.8) as estimated by quantitative PCR (linear mixed effect modeling, p value < 0.001; **Figure 2D**), possibly caused by MccI47 accumulation during the growth phase. This suggests that even though we increased the plasmid size 2.6-fold, pMut2-I47 is retained stably long-term without the need of selection. We then confirmed *EcN*^*ΔH ΔM*^*-I47* killing activity using static inhibition assays. *EcN*^*ΔH ΔM*^*-I47* produced a zone of inhibition of more than 2.3 cm in diameter against CR *K. pneumoniae* (BAA 1705) irrespective of the addition of ampicillin for antibiotic plasmid retention (**Figure 2E**). Simultaneously, the two respective controls *EcN* (1) and *EcN*^*ΔH ΔM*^ (2) were unable to produce any halo of inhibition. Taken together, these data provide the evidence for a novel *EcN* strain engineered to constitutively produce MccI47 from a self-retaining plasmid *in vitro*, which is suitable for application in a murine model to challenge CR *K. pneumoniae* colonization.

**Figure 2:**
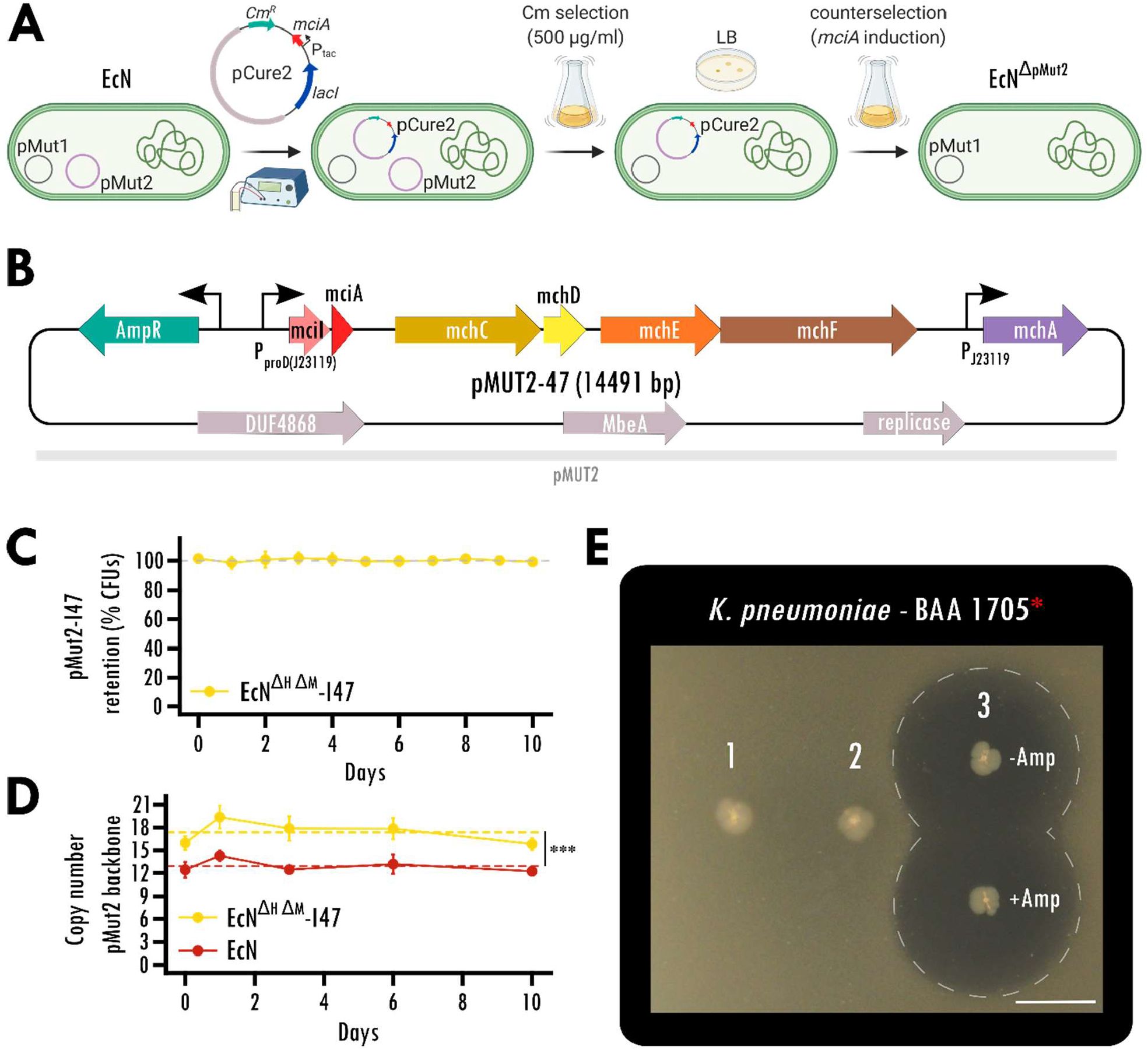
Stable MccI47 production from a native, self-retaining *EcN* plasmid. (A) Curing of the native *EcN* plasmid pMut2 *via* utilizing inducible MccI47 expression as a counterselection tool. Counterselection plasmid pCure2-I47 with pMut2 as backbone was introduced into *EcN* by electroporation and chloramphenicol (Cm) supplementation. High concentrations of Cm led to complete displacement of pMut2 in the cells. Without further antibiotic selection, *mciA* expression was induced, leading to selection against pCure2-I47, since the immunity peptide *mciI* was omitted in the plasmid design. (B) Plasmid map of pMut2-I47, an MccI47-producing plasmid with pMut2 as backbone vector. (C) Serial passage experiment to assess pMut2-I47 plasmid retention in *EcN*^Δ*H* Δ*M*^*-I47*. Cells were initially grown with the supplementation of ampicillin and then subjected to daily subculturing for ten days without selection pressure, resulting in a total of approximately 200 bacterial generations. n=5. (D) Assessment of plasmid copy number of pMut2 and the pMut2-based pMut2-I47 using quantitative PCR. Note that *EcN*^Δ*H* Δ*M*^*-I47* cells harboring pMut2-I47 exhibit a significantly higher copy number mean compared to *EcN* harboring native pMut2. Additionally, both plasmids are stably retained for the ten days of the serial passage experiment. ***: p < 0.001. n=5. (E) Static inhibition assay comparing inhibitory activity of (1) *EcN*, (2) *EcN*^*ΔH ΔM*^, and (3) *EcN*^*ΔH ΔM*^*-I47*, harboring pMut2-I47, against carbapenem-resistant *Klebsiella pneumoniae*. Inhibitory activity of *EcN*^*ΔH ΔM*^*-I47* is independent ampicillin (Amp) for plasmid selection. Scale bar: 1 cm.

To test whether *EcN*^*ΔH ΔM*^*-I47* can reduce CR *K. pneumoniae* colonization *in vivo*, we performed a decolonization experiment in a specific-pathogen-free mouse model[38-40]. Briefly, we treated C57BL/6J mice with ampicillin for seven days. At day five, mice were administered 10^8^ cells of CR *K. pneumoniae* (BAA 1705), for which we had demonstrated MccI47-mediated inhibitory effects in **Figures 1B, C, Table 1**, and **Figure 2D**. One week after CR *K. pneumoniae* application, each mouse was orally administered 10^8^ cells of *EcN, EcN*^*ΔH ΔM*^ or *EcN*^*ΔH ΔM*^-I47 daily for seven days (**Figure 3A**). Fecal samples were collected throughout the experiment and plated to quantify abundances of CR *K. pneumoniae* and the respective *EcN* treatment strains. Linear regression analysis of the log fold change in CR *K. pneumoniae* colonization before and after treatment as a function of the mean abundance of each treatment strain (*i*.*e*., *EcN, EcN*^*ΔH ΔM*^, or *EcN*^*ΔH ΔM*^*-I47*), quantified as shedding, revealed that *EcN* does not seem to affect *K. pneumoniae* colonization *in vivo* (regression coefficient = -0.008, p = 0.994). However, we found that increasing abundance of *EcN*^*ΔH ΔM*^ resulted in higher abundance of *K. pneumoniae* (regression coefficient = 1.856, p = 0.029), underlining the previously described role of class IIb microcins in niche competition. Strikingly, the effect of *EcN*^*ΔH ΔM*^*-I47* on *K. pneumoniae* was significantly different compared to its control *EcN*^*ΔH ΔM*^ (p = 0.006) as an increase in abundance of *EcN*^*ΔH ΔM*^*-I47* caused a reduction in CR *K. pneumoniae* colonization (regression coefficient = -2.504, p = 0.047) (**Figure 3B**) in the murine model. To determine whether the treatments resulted in alterations of the resident microbiota, we analyzed the relative bacterial abundances before and after treatment through 16S rRNA profiling. Principal component analysis of pre and post treatment groups revealed significant community changes for all samples (PERMANOVA, p < 0.0001) likely caused by the time passed since the antibiotic treatment and the recovery of the resident microbiota (**Figure 3C**). However, we could not find any live biotherapeutic product-dependent changes in the community caused by the different *EcN* strains (PERMANOVA, p = 0.81). Differential analysis using DESeq2 resulted in only four significantly different bacterial species across treatments (*Adlercreutzia equolifaciens, Anaeroplasma abactoclasticum, Harryflintia acetispora, Ruminiclostridium cellulolyticum*). However, this may be due to the use of sensitive statistical testing, since none of them displays consistent signal over most samples across the treatment groups (**Figure S5**). Therefore, we can conclude that in contrast to commonly prescribed antibiotics, treatment with the engineered strain has minimal effects on the GI microbiota, with the addition that our engineered biotherapeutic strain *EcN*^*ΔH ΔM*^*-I47* selectively targets CR *K. pneumoniae*. Taken together these data demonstrate that the oral delivery of an engineered probiotic *E. coli* strain overproducing MccI47 while passing through the GI tract is a viable approach to reduce CR *K. pneumoniae* colonization *in vivo*.

**Figure 3:**
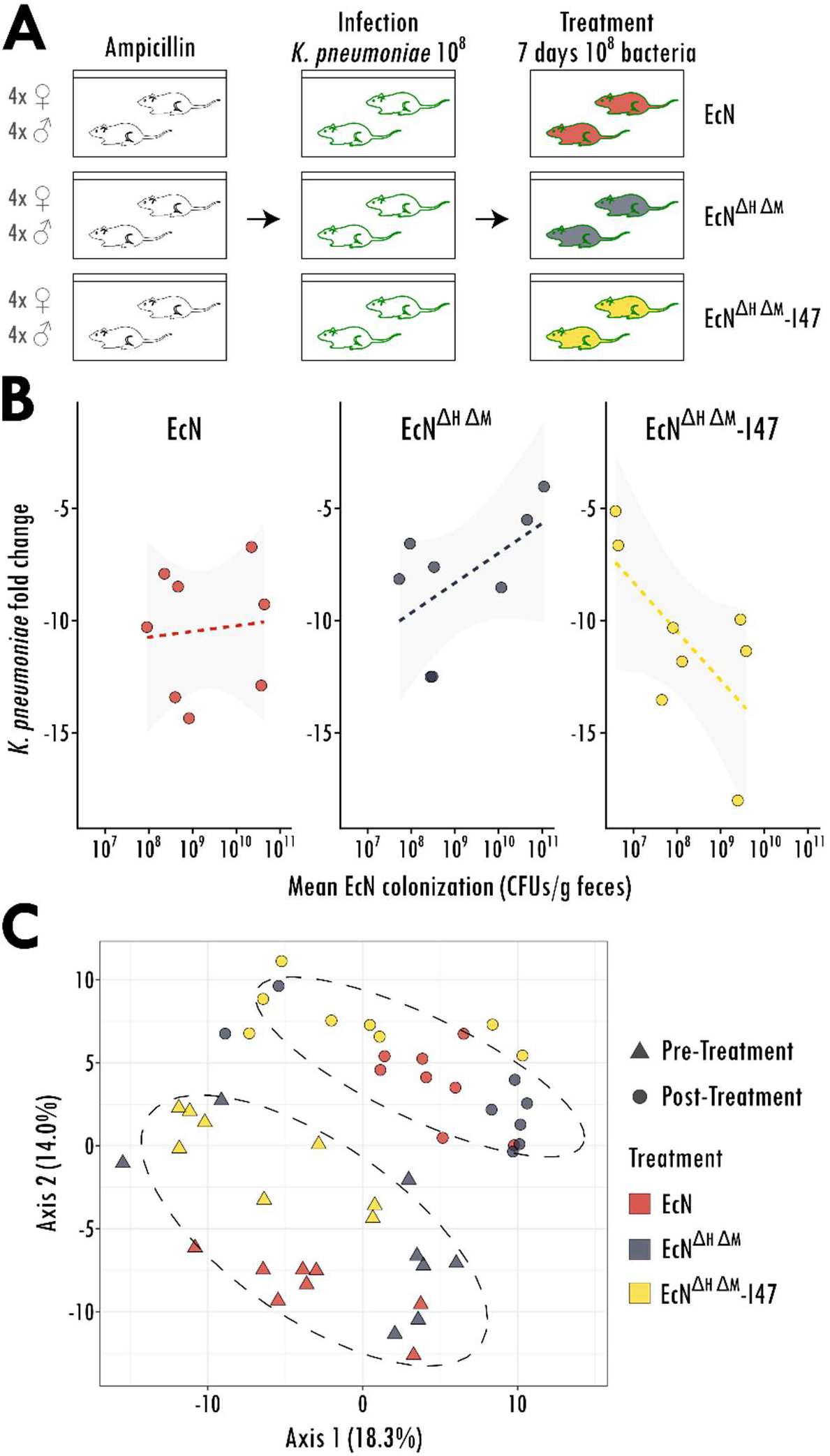
Engineered probiotic expressing MccI47 is effectively reducing colonization of carbapenem-resistant *Klebsiella pneumoniae in vivo*. (A) Schematic representation of the experimental setup. Male and female mice were treated with ampicillin in the drinking water for seven days and infected with 10^8^ cells of carbapenem-resistant *K. pneumoniae* (BAA-1705) on day five of the experiment. Starting on day twelve, mice were treated daily with 10^8^ cells of the different *EcN* strains (*EcN, EcN*^*ΔH ΔM*^ and *EcN*^*ΔH ΔM*^*-I47*) for seven days by oral administration. (B) Linear mixed effect modeling regression was used to assess the log fold change in *K. pneumoniae* (BAA 1705) colonization before and after treatment as a function of the mean shedding of each respective *EcN* treatment strain (*EcN, EcN*^*ΔH ΔM*^ or *EcN*^*ΔH ΔM*^*-I47*). (n=8). (C) Principal component analysis (PCoA) of the bacterial communities of every mouse before and after treatment. Note that the experimental time introduces significant differences between the samples pre and post treatment, however, no treatment-specific differences between *EcN, EcN*^*ΔH ΔM*^ or *EcN*^*ΔH ΔM*^*-I47* were observed. Ellipses are grouping samples according to time point pre and post treatment. (n=8).

## Discussion

Under the global societal challenge of moving towards more personalized medicine, biomedical research is currently exploring the use of engineered bacteria as vehicles for the targeted administration of therapeutic compounds at the location of disease and with the goal of reducing systemic side effects[33, 41, 42]. This is particularly relevant when the therapeutic objective is the delivery of protein-based agents (*i*.*e*., AMPs) to the GI tract as host digestive enzymes likely inactivate them before they reach their site of action[43]. Furthermore, embedding the proteins into a lipid matrix within enteric capsules to prevent host-mediated degradation[44], needs to account for the agent’s dilution effect that naturally occurs downstream of the point of capsule’s opening. These issues are circumvented with the use of live, engineered bacteria as they can be designed to ensure continuous production of the therapeutic compound while passing through the GI tract[45].

Current antibiotics are often applied systemically to treat bacterial pathogens, yet increasingly fail to clear infections caused by MDR organisms[46]. Additionally, these drugs also cause profound gastrointestinal dysbiosis of the protective microbiota, leaving treated individuals with often dangerous secondary outcomes[47]. Selective decolonization of pathogenic bacteria as a way to minimize their translocation and horizontal transmission to other patients is currently being explored in the pharmaceutical industry by assembling consortia of isolated bacteria that display decolonizing properties against certain species, including *K. pneumoniae*[48]. However, to our knowledge there is no biotherapeutic yet that has reach stage for human trials.

Inspired by recent research on the role of siderophore microcins in modulating the dynamics of gastrointestinal bacteria[49], we here provide the first every characterization of the to-date only putative class IIb microcin MccI47. We show that MccI47 is a very potent antimicrobial against multiple MDR *Enterobacteriaceae* strains and displays strong inhibitory effect against CR *K. pneumoniae*. Based on this we then develop a rationally engineered biotherapeutic *E. coli* Nissle 1917 that is capable of MccI47 overproduction. Compared to previous studies that either rely on genomic integration [32] (which would lead to a single copy of the operon and thus suboptimal production) or rely on the use of plasmids that require a strong selective pressure for their maintenance[50], we devised a novel constitutive expression system from a stably-retained multicopy plasmid that allows for high MccI47 output, while rendering provisions for plasmid selection unnecessary. Finally, we demonstrate clinical relevance of our engineered strain in a in vivo model CR *K. pneumoniae* colonization by showing a significant reduction in CR *K. pneumoniae* intestinal abundance after seven days of daily oral probiotic administration, with no major effect on the resident microbiota.

## Conclusions

This study provides the first ever characterization of a to-date only putative microcin with potent inhibitory effect against CR *K. pneumoniae*. To show its effect *in vivo* we develop a novel strain of the probiotic EcN that can produce this CR *K. pneumoniae-*killing microcin from a stably retained engineered native plasmid without the need of any selection. We provide the first demonstration of microcin-mediated modulation of CR *K. pneumoniae* dynamics *in vivo*. Overall, this study serves as the foundational step towards the use of engineered probiotics and engineered live biotherapeutic products aimed at the selective removal of MDR pathogens from the GI tract. We believe that this study will open the door to investigations into the optimization, scale-up and manufacturing of these next generation therapeutic agents and eventually human trials.

## Methods

### Bacterial strains and plasmids

This study used the *Escherichia coli* strains NEB10β (*New England Biolabs, Ipswich, MA*), Nissle 1917 as well as all strains listed in Table 1. Strains in Table 1 were purchased from ATCC (*Manassas, VA*). All plasmids in this study have been transformed into cells using electroporation with the Bio-Rad Micropulser™ (*Bio-Rad Laboratories, Hercules, CA*) and were created using Gibson Assembly[51] using the Gibson Assembly Master Mix (*New England Biolabs, Ipswich, MA*) and custom DNA oligonucleotides purchased from Integrated DNA Technologies (*Coralville, IA*). For pBBAD-H47 and pBBAD-I47, four fragments were amplified by PCR and assembled: (1) linearized pUC19, (2) *araC* and P_BAD_ from pTARA[52] (Addgene #39491), (3) the microcin and immunity genes for MccH47 (*mchXIB*) or MccI47 (*mciIA*) originating from pEX2000[53] as well as (4) the genes *mchCDEFA* originating from pPP2000[20]. Plasmid pHMT-H47 has been described and used in[20]. For plasmid pHMT-I47 a total of seven fragments was assembled: (1) linearized pUC19, (2) chloramphenicol resistance cassette from pTARA[52] (Addgene #39491), (3) *lacI* and *tac* promoter from pMAL-c5X (*New England Biolabs, Ipswich, MA*), (4) MBP, amplified using primers to add a 6× Histidine N-terminal tag, from pMAL-c5X, (5) *mciA* from pEX2000[53], (6) *mciI* from pEX2000, and (7) *mchCDEFA* from pPP2000. To cure the native pMut2 plasmid from *E. coli* Nissle 1917, pCure2-I47 was assembled from four fragment: (1) linearized pMut2, (2) chloramphenicol resistance cassette from pTARA[52] (Addgene #39491), (3) *mciA* from pEX2000[53], (4) *lacI* and *tac* promoter from pMAL-c5X (*New England Biolabs, Ipswich, MA*). The modified replacement plasmid for pMut2, pMut2-I47, was created from four fragments: (1) linearized pMut2, (2) ampicillin resistance cassette from pUC19, (3) insulated promoter proD[37], where the promoter region was replaced with the strong constitutive promoter J23119 (BBa_J23119), (4) *mciIA* to *mchCDEFA* from pBBAD-I47. All plasmid sequences and maps have been deposited as .dna files at: https://gitlab.com/vanni-bucci/2021_MccI47_paper. Chromosomal modifications were obtained through lambda Red recombination using pKD46 and FLP-FRT recombination using pCP20 as described by Datsenko and Wanner[54]. Briefly, a kanamycin resistance cassette flanked by flippase recognition target (FRT) sites was amplified by PCR from pKD4 and then transformed into the respective electrocompetent *EcN* strain harboring plasmid pKD46. For strain *EcN*^*ΔH ΔM*^ the kanamycin resistance was removed after the knockout of *mchIB* by inducing the flippase from pKD20, before it was reintroduced for the *mcmIA* knockout. All modifications were confirmed using Sanger sequencing (**Figure S3**).

### Plasmid curing and assessment of retention

To cure the native plasmid pMut2 from *E. coli* Nissle 1917, the inducible plasmid pCure2-I47 was created for counterselection. After transformation of pCure2-I47 into *E. coli* Nissle 1917, plasmid presence was confirmed by selective plating on chloramphenicol (25 µg/ml) and PCR. Positive clones were individually grown over night at 37°C under strong antibiotic selection (500 µg/ml) in liquid LB medium to skew plasmid competition towards pCure2-I47 and to displace pMut2. Resulting cultures were plated on LB agar and the loss of pMut2 was confirmed by PCR (**Table S1**). Resulting clones were again individually grown over night at 37°C in LB with the addition of 0.5 mM IPTG. Since pCure2-I47 expresses the MccI47 toxin (*mciA*) without the immunity peptide (*mciI*) under IPTG induction, cells harboring pCure2-I47 are killed in response to the addition of IPTG and therefore forced to lose the plasmid to survive. Resulting cultures were plated on LB agar and loss of pMut2 and pCure2-I47 was confirmed by PCR (**Figure S2B**). Subsequently, pMut2-I47 was transformed into cured cells and retention without antibiotic selection was assessed by a serial passage experiment. Five independent replicates of *EcN*^*ΔH ΔM*^*-I47* were initially grown under ampicillin selection (100 µg/ml) overnight at 37°C in liquid LB medium (to day 0) and then subcultured 10^−6^ daily for ten days (until day 10), allowing for the growth of approximately 20 bacterial generations per passage. Each biological replicate was plated daily on LB agar using three technical replicates with and without ampicillin (100 µg/ml) to quantify the percentage of CFUs retaining the plasmid over time. To test for the effect of time on the number of CFUs of *EcN*^*ΔH ΔM*^*-I47* retaining pMut2-I47 we ran linear mixed effect modeling in ‘R’ using the *lmer* from the lmerTest package. Specifically, we fit models predicting the percentage of CFUs having the pMut2-I47 plasmid as a function of time and using replicate ID as random effect. To assess the average plasmid’s copy number per bacterium, cells contained in 1 ml from each daily passed culture were harvested and frozen at -80°C until qPCR. To compare pMut2-I47 copy number with the copy number of the native pMut2, *EcN* was also subjected to the same serial passage assay and related procedures as control. Plasmid copy numbers have been assessed using qPCR by comparing the signal from the plasmid pMut2 to that from the chromosomal gene DNA gyrase subunit B (**Table S1**)[55]. The frozen bacterial pellet was resuspended in 1 ml of sterile PBS, incubated for 5 min at 95°C and used as template in the qPCR reaction in a final 10^−4^ dilution with the iTaq Universal SYBR Green Supermix (*Bio-Rad Laboratories, Hercules, CA*). The qPCR was run in a StepOnePlus Real-Time PCR System (*Applied Biosystems, Waltham, MA*) and plasmid copy number was calculated using the ΔΔCt, assuming an efficiency of 100%. To test for differences in the number of copies of pMut2-I47 (carried by *EcN*^*ΔH ΔM*^*-I47*) versus the number of copies of pMut2 (carried by *EcN*) and the effect of time, we ran linear mixed effect modeling in ‘R’ using the *lmer* from the lmerTest package. Here we fit models where the copy number was predicted as a function of the strain (*i*.*e*., *EcN*^*ΔH ΔM*^*-I47* vs. *EcN*), time and the interaction of the two and using replicate ID as random effect.

### MccI47 purification

The MBP-MccI47 fusion protein was expressed from pHMT-I47 and purified as described in[20]. Briefly, overnight cultures of the *E. coli* NEB10β strain harboring pHMT-I47 were grown in LB under antibiotic selection (100 µg/ml ampicillin, 25 µg/ml chloramphenicol). The culture was diluted 1:50 in prewarmed LB medium and incubated at 200 rpm and 37°C. After 90 min 2,2-dipyridyl was added to a final concentration of 0.2 mM. After two more hours (approximate optical density at 600nm OD_600_=0.2), MccI47 production was induced with 0.5 mM IPTG for five hours. The cells were then harvested by centrifugation, resuspended in column buffer (200 mM NaCl, 20 mM Tris-HCl, pH 7.5) and frozen until purification at -20°C. The cells were slowly thawed in cold water, sonicated, and centrifuged at 4°C with 15,000xg for 10 min to remove the debris. The lysate was then passed through an amylose resin (*New England Biolabs, Ipswich, MA*) column and finally the MBP-MccI47 fusion proteins were eluted into 30 ml using 10mM maltose in the column buffer. The MBP-MccI47-containing solution was concentrated using MilliporeSigma (*Burlington, MA*) MWCO 10,000 filters and then digested with 10 µl Tobacco etch virus nuclear-inclusion-a endopeptidase (TEV) (*New England Biolabs, Ipswich, MA*) overnight at 4°C. The next day the solution was brought to room temperature, another 5 µl TEV were added, and the digestion was incubated for additional two hours, resulting in free MccI47 in the solution. The histidine-tagged TEV and MBP were extracted from the solution using a Ni-NTA agarose resin (*Qiagen, Hilden, Germany*) as described in[20].

### Plate inhibition assays

Solid media inhibition assays were performed as described previously[20, 21]. Briefly, individual bacterial colonies of the microcin-producing strains (MccH47 or MccI47) were picked with a sterile pipet tip and stabbed into the solid LB agar. Iron-limited conditions during the growth phase were created by supplementing the media with 0.2 mM 2,2-dipyridyl. For the inducible constructs pBBAD-MccH47 and pBBAD-MccI47 the media was supplemented with 100 µg/mL ampicillin for plasmid retention and 0.4% L-arabinose for microcin production. For the self-retaining plasmid pMut2-MccI47 no inducing agent was necessary. Here, were indicated, ampicillin was locally added to the media by drying a drop of sterile water containing 500 µg ampicillin before adding the colony. All plates were incubated for 48 h and the bacteria were inactivated with chlorophorm vapors and 10 min of ultraviolet light. For the overlay, target bacteria were diluted from an overnight culture (*E. coli* BAA 196 – 1:200; *K. pneumoniae* BAA 1705 – 1:1000) in 3 ml LB with 0.2 mM 2,2-dipyridyl and 100 µg/mL ampicillin. Then molten agar was added to the media to a final concentration of 0.75%. 3.5 ml of the soft agar medium was then evenly spread on the plate, cooled and the plate was incubated for 16 h at 37°C before imaging.

### Minimum inhibitory concentration (MIC) assays

The MIC assays were carried out in sterile 96-well round bottom microplates (*CELLTREAT Scientific Products, Pepperell, MA*) and were set up with the following media. The first row was filled with 20 µl 2x LB with 0.4 mM 2,2’-dipyridyl and 20 µl of purified MccI47 in amylose resin elution buffer (200 mM NaCl, 20 mM Tris-HCl, 10 mM maltose, pH 7.5), resulting in the highest MccI47 concentration for the assay in 1x LB, 0.2 mM 2,2’-dipyridyl, 0.5× amylose resin elution buffer. All other wells were loaded with 20 µl 1x LB, 0.2 mM 2,2’-dipyridyl, 0.5× amylose resin elution buffer and two-fold serial dilution were performed across the plate. Target bacteria were grown over night in standard LB at 200 rpm at 37°C, and added to a final dilution of 10^−4^ into the individual wells. The plates were incubated in the dark at room temperature with gentle agitation and MICs were determined as the lowest concentration for which no growth was observed after 24 h. All reported MIC values represent the median of at least three biological replicates from independent MccI47 purifications.

### Animal study

The mouse experiment was carried out under an institutionally approved Institutional Animal Care and Use Committee (IACUC) protocol. Per arm eight C57BL/6J mice (four male, four female) at seven weeks of age were housed in same-sex pairs in four cages under specific pathogen-free conditions. Each mouse was individually marked, so they could be traced throughout the entire experiment. To achieve robust gastrointestinal CR *Klebsiella pneumoniae* (strain BAA 1705) engraftment, all mice were treated with 0.5 g/l ampicillin in the drinking water for seven days. On day five of the experiment (during ampicillin treatment) mice were deprived of food and water for four hours and then given 10^8^ cells of CR *K. pneumoniae* (strain BAA 1705) in 20 µl phosphate buffered saline (PBS) by pipet. Starting on day twelve of the experiment, the three arms were given daily 10^8^ cells for seven days of either (i) *EcN*, (ii) *EcN* with a knockout of the genes *mchIB* (MccH47) and *mcmIA* (MccM) to prevent interference of the other class IIb microcins in the *EcN* genome (*EcN*^ΔH ΔM^), or (iii) *EcN* with the same gene knockouts and constitutive MccI47-production from a modified version of the native plasmid pMut2 (*EcN*^*ΔH ΔM*^-*I47*). For administration the mice were again deprived of food and water for four hours and then given the cells orally by pipet in 20 µl PBS. The last day of the treatment was day 18. Fecal samples were taken on day twelve before first administration, day 13, 15, and day 19 to assess the colonization of the treatment strains as well as bacterial shedding of CR *K. pneumoniae* (strain BAA 1705). For sampling, mice were placed into sterile isolation containers and at least two fecal samples (approximately 0.03 g each) were collected into sterile two 1.5 ml microcentrifuge tubes. One sample was snapfrozen and stored at -80°C until DNA extraction and 16S rRNA amplicon sequencing, the other one was weighed, resuspended in 300 µl sterile PBS and kept on ice until plating. To determine the bacterial shedding in CFU/g feces, the samples were shortly spun in a minicentrifuge to pellet larger particles and plated in 10-fold serial dilutions from the supernatant on respective antibiotic plates and incubated at 37°C for 16 h, which resulted in a detection limit at 10^2^CFU/g feces. To test for the effect of treatment on CR *K. pneumoniae* colonization we ran linear regression models. Specifically, we use linear-mixed effect modeling to predict the log fold change in CFUs of CR *K. pneumoniae* between after and before biotherapeutic treatment as a function of the mean colonization of each probiotic strain, the strain type, and the interaction of the two as fixed effects and using the cage ID as random effect. The model was fitted to the data using the *lmer* function from the lmerTest package in ‘R’ and run three times corresponding to setting each strain as baseline to determine significance and magnitude of the strain-specific coefficients on CR *K. pneumoniae* fold change. Significance and magnitude of interaction coefficients were inspected to determine differences among the treatment type. Data and ‘R’ code to perform statistics have been deposited at: https://gitlab.com/vanni-bucci/2021_MccI47_paper.

### Relative bacterial abundance analyses

DNA was extracted from the frozen fecal pellets with the DNeasy Powersoil Pro Kit by Qiagen (*Hilden, Germany*) according to the manufacturer’s protocol. The bacterial 16S rRNA gene (variable regions V3 to V4) was subjected to PCR amplification using the using the universal 341F and 806R barcoded primers for Illumina sequencing. Using the SequalPrep Normalization kit, the products were pooled into sequencing libraries in equimolar amounts and sequenced on the Illumina MiSeq platform using v3 chemistry for 2 × 300 bp reads. The forward and reverse amplicon sequencing reads were dereplicated and sequences were inferred using dada2[56]. Differential microbiome analysis and visualization was performed in ‘R’ using DESeq2[57]. All ‘R’ code has been deposited at: https://gitlab.com/vanni-bucci/2021_MccI47_paper. Raw microbiome data have been deposited on the European Sequencing Archive (ENA) accession number PRJEB48537.

## Availability of data and materials

All data and code to create the figures of this study as well as DNA files for genetic engineering work are available at https://gitlab.com/vanni-bucci/2021_MccI47_paper. Raw reads for the 16S rRNA amplicon sequencing are deposited on the European Sequencing Archive (ENA) under accession number PRJEB48537. Plasmids and strains are available from the authors upon request.

## Competing interest

V.B. receives support from a sponsored research agreement from Vedanta Biosciences, Inc.

## Funding

This work was supported by the CDRMP PRMP W81XWH2020013 to V.B. and M.W.S, by the UMass Office of the President Science and Technology Award to V.B., M.W.S, and B.A.M. and by the Deutsche Forschungsgemeinschaft (DFG) project MO 4092/1-1 to B.M.M.

## Author Contribution

B.M.M, J.D.P, V.B, M.W.S and B.A.M conceptualized the study. B.M.M, J.D.P, V.B designed the experiments. B.M.M and J.D.P performed initial engineering and testing of MccI47-MBP expressing strains. B.M.M and H.D performed purification and *in vitro* minimum inhibitory concentration assays. B.M.M conceptualized and performed EcN stable engineering and testing. B.M.M and R.ML performed *in vivo* experiments. C.B performed microbiome sequencing. S.K.B performed microbiome analytics. V.B and B.M.M performed statistical testing and data interpretation. M.W.S and B.A.M supported with data interpretation and with microbiological and infectious disease expertise. B.M.M and V.B wrote the manuscript with input of all authors.

## Acknowledgements

We appreciate the help and microbiological expertise by Kenan Murphy. Figure 2A was created with BioRender.com. Figure S3 was created using Easyfig[58].

## Notes

https://gitlab.com/vanni-bucci/2021_MccI47_paper

