## Supplementary Materials for "Microcin MccI47 selectively inhibits enteric bacteria and reduces carbapenem-resistant *Klebsiella pneumoniae* colonization *in vivo* when administered *via* an engineered live biotherapeutic"

**Affiliations:**

**Supplementary Table 1**: Primers used for plasmid verification in Supplementary Figure 2B and quantitative PCR in Figure 2D.

| Primer name | Sequence | Target | Fragment size |
|---|---|---|---|
| pMut1_MobA_F<br>pMut1_ORF4_R | GTGCCCTGTTATCCAGGCTTATGG<br>AGGTTGAAGGTCTCAGAGAATGAGAC | pMut1 | 1208 bp |
| pMut2_ORF2_F<br>pMut2_R | ATGTTAATCTGCTATTTGAATAGTCGAGTACGC<br>GCTCGTCATCGATCCGAATATTAATCG | pMut2 | 1296 bp |
| pMut2_ORF2_F<br>pCure2_lacI_R | ATGTTAATCTGCTATTTGAATAGTCGAGTACGC<br>CGACATCGTATAACGTTACTGGTTTCAC | pCure2-I47 | 1493 bp |
| pMut2_ORF2_F<br>pMut2_AmpR_R | ATGTTAATCTGCTATTTGAATAGTCGAGTACGC<br>TCTACACGACGGGGAGTCAGG | pMut2-I47 | 244 bp |
| qPCR_gyrB_F<br>qPCR_gyrB_R | CATGGAGCGTCGTTATCCGA<br>CTGCCGTGCTGTTCTTTGTC | gyrB | 147 bp |
| qPCR_pMut2_F<br>qPCR_pMut2_R | CAAAGCCCCGAAATCATGCTC<br>CGGAGAAGTACGGCTTGTGG | pMut2 | 186 bp |

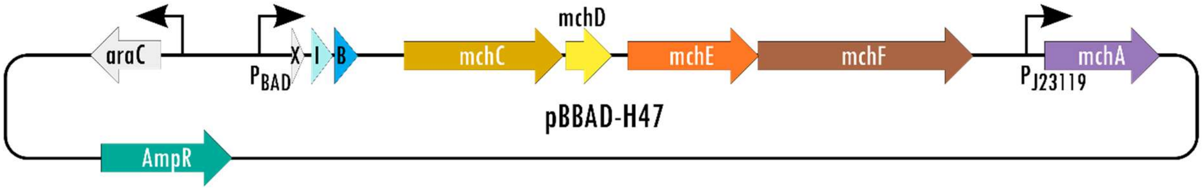


**Supplementary Figure 1**: Plasmid map of pBBAD-H47, a pUC19-based plasmid producing MccH47-MGE from *mchB*, the immunity peptide *mchI* under an L-arabinose inducible $P_{BAD}$ promoter as well as the genes needed for post-translational modification *mchCDEFA*. AmpR = ampicillin resistance, X = *mchX*, I = *mchI*, B = *mchB.*

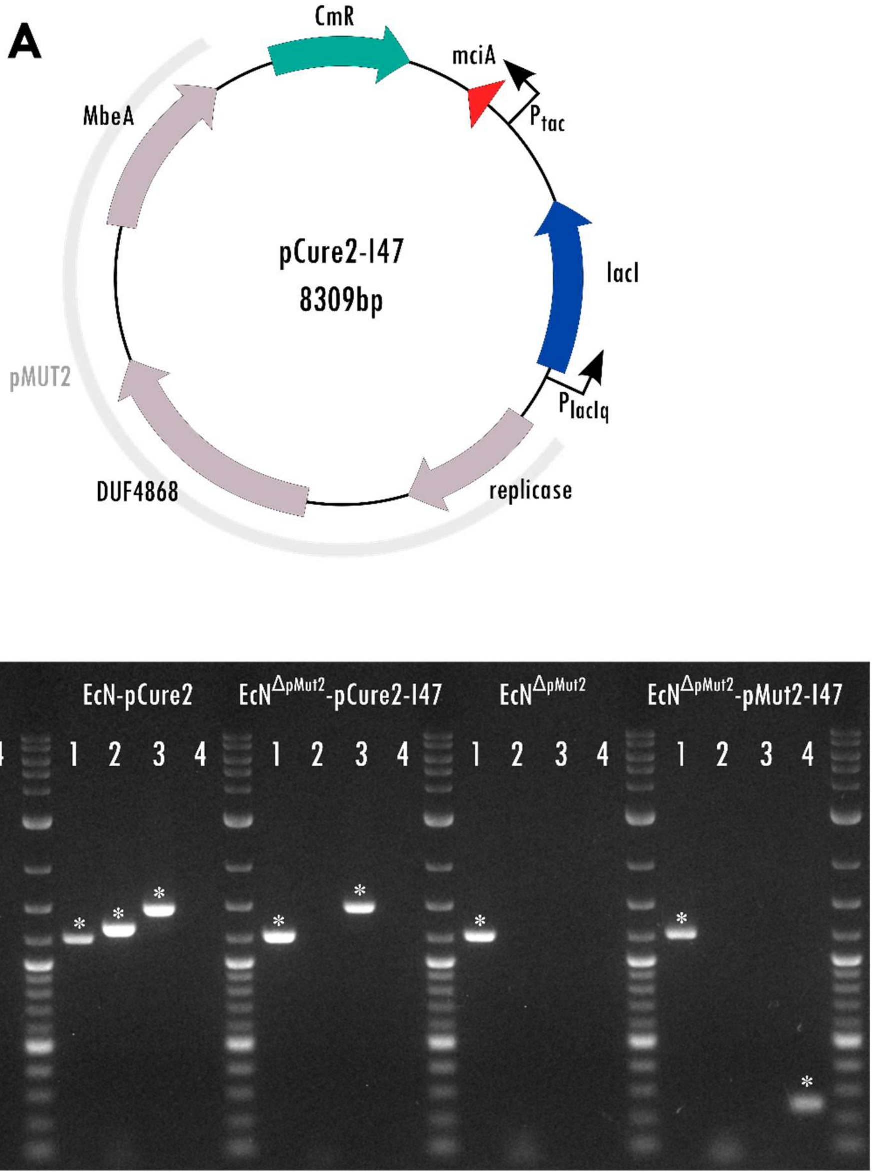


**Supplementary Figure 2**: (A) Plasmid map of pCure2-I47. Grey shaded area indicates the pMut2 backbone. CmR = chloramphenicol resistance. (B) Gel electrophoresis of PCR products for *EcN* strains created during the process pMut2 curing and transformation with pMut2-I47. (1) Primers against pMut1 (1208 bp), (2) primers against the insert-free pMut2 backbone (1296 bp), (3) primers against pCure2-I47 (1493 bp), (4) primers against pMut2-I47 (244 bp). Asterisks indicate specific fragments amplified by PCR. 1kb Plus DNA Ladder (*New England Biolabs, Ipswich, MA*).

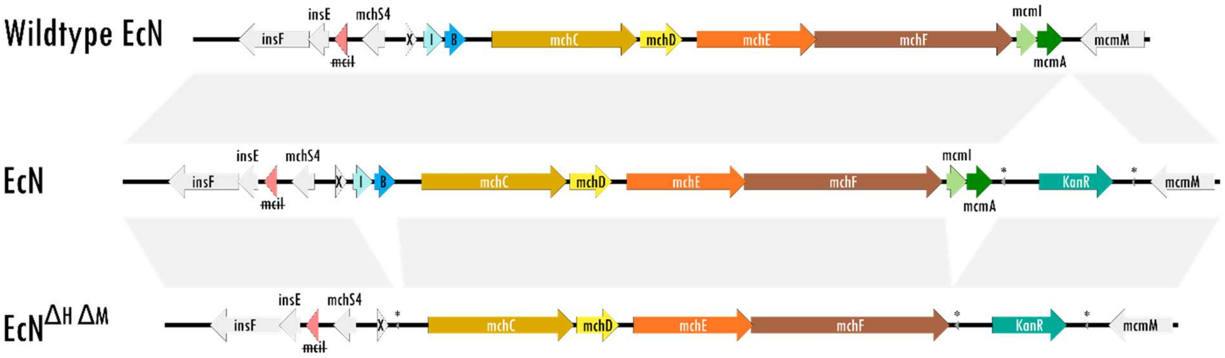


**Supplementary Figure 3**: Genetic organization of the *E. coli* Nissle 1917 class IIb microcin gene cluster. Wildtype *EcN* (top) harbors genes for production of the microcins MccH47 (*mchB*) and microcin MccM (*mcmA*) including their corresponding immunity genes *mchI* and *mcmI.* To allow selective colony counting, a kanamycin resistance cassette was introduced into the *EcN* genome and the resulting strain served as the *EcN* control in this study (center, *EcN*). A MccH47/MccM knockout strain (bottom, $EcN^{\Delta H\,\Delta M}$) was generated to exclude the interference of native *EcN* microcin production, when assessing the potency of MccI47 against *K. pneumoniae in vitro* and *in vivo*. This strain served as the knockout control and the basis for the strain harboring the MccI47-producing plasmid pMut2-I47 ($EcN^{\Delta H\,\Delta M}$*-I47*). Shaded areas indicate sequence similarity between the depicted gene clusters. X = *mchX*, I = *mchI*, B = *mchB,* ~~mciI~~ = truncated mciI, * = flippase recognition target (FRT) site, KanR = kanamycin resistance.

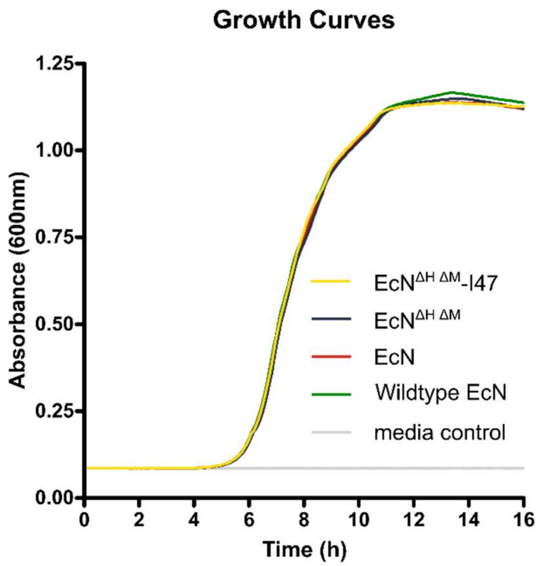


**Supplementary Figure 44:** Growth curves of different *EcN* strains. Note that none of the genetic modifications had an impact on the growth rate. Wildtype *EcN* represents the native strain, while all other strains were created for this study (see Supplementary Figure 3). *EcN* and $EcN^{\Delta H\,\Delta M}$ serve as the study controls for $EcN^{\Delta H\,\Delta M}$-I47, which is expressing MccI47 from pMut2-I47. n = 8.

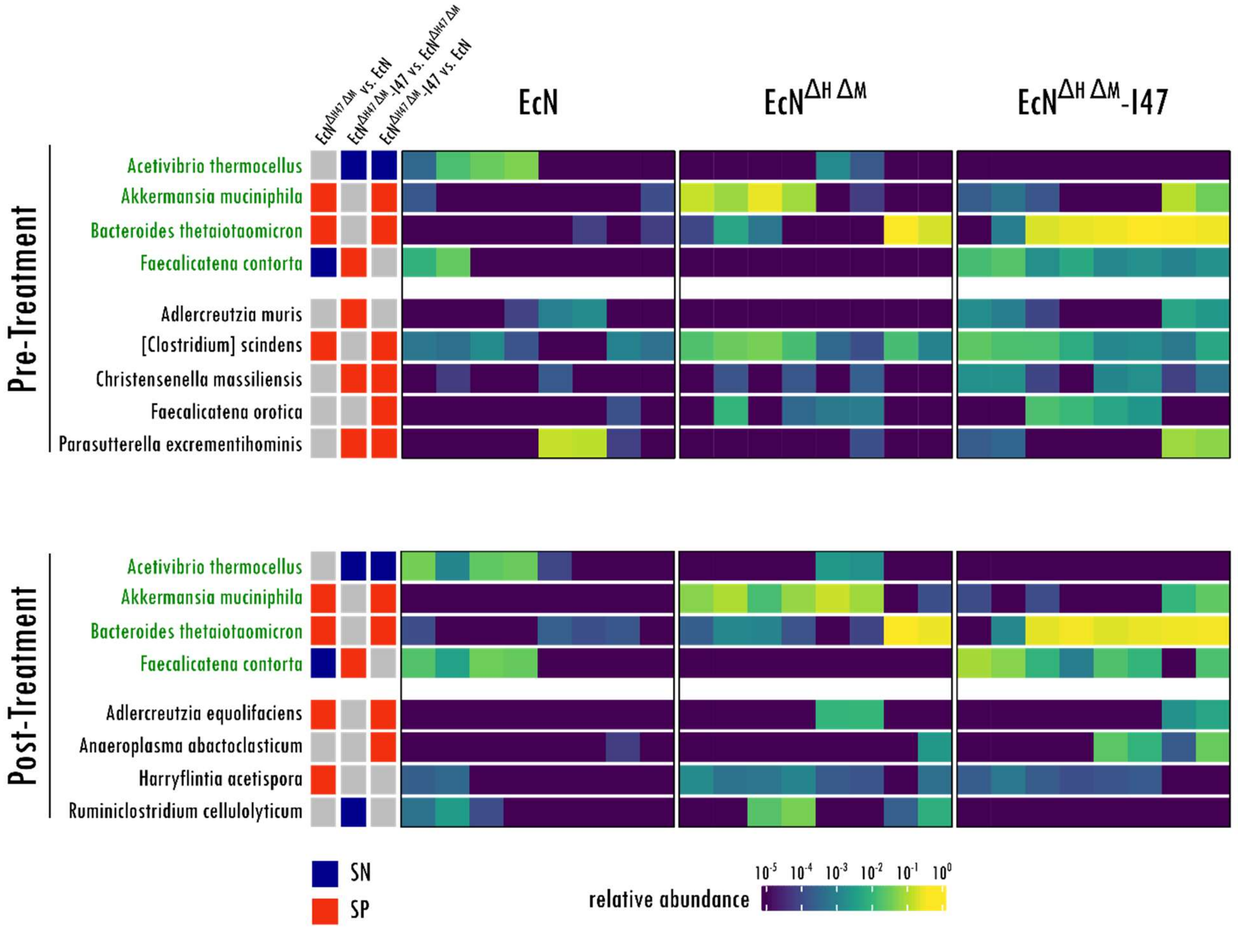


**Supplementary Figure 5:** Heatmap of sensitive differential bacterial species analysis before and after treatment. Note that four of the differential species are significantly different between groups pre and post treatment with *EcN*, *EcN$^{\Delta H \Delta M}$* or *EcN$^{\Delta H \Delta M}$*-I47 (green) and therefore do not reflect a change induced by any of the treatment strains. SN = significant negative change, SP=significant positive change.
